# HB21/40/53 promote inflorescence arrest through ABA accumulation at the end of flowering

**DOI:** 10.1101/2023.04.20.537726

**Authors:** Verónica Sánchez-Gerschon, Cristina Ferrándiz, Vicente Balanzà

## Abstract

Flowers are produced by the activity of the inflorescence meristem after the floral transition. In plants with indeterminate inflorescences, as Arabidopsis, the final number of flowers produced by the inflorescence meristem will depend on two main factors, the rate of flower production by the meristem and the duration of the phase of inflorescence meristem activity. The end of flowering, understood as the moment when the inflorescence stops the production of new flowers, is associated with the meristem proliferative arrest. At this time point, the meristem ceases to initiate new floral primordia and the unpollinated flowers already formed arrest their development.

It has been known for a long time that fruit/seed production induces inflorescence meristem arrest, but the mechanisms controlling this process were elusive. During the last years, the regulation of the end of flowering has started to be elucidated in Arabidopsis. The meristem arrest at the end of flowering is controlled at the genetic level by the FRUITFULL-APETALA2 (FUL-AP2) pathway, that modulates meristem activity. The meristem arrest has been also shown to be controlled at the hormonal level. It has been proposed that auxin could mediate the fruit/seed effect to the meristem. Cytokinins regulation and response have been also proposed as important factors controlling the meristem activity at the end of flowering. Finally, it has been also described that arrested meristems at the end of flowering resembles dormant meristem at the transcriptomic level.

Previously, we have shown that the FUL-AP2 pathway controls the expression of the homeodomain leucine zipper transcription factor *HOMEOBOX PROTEIN 21* (*HB21*), a gene involved in the establishment of bud axillary dormancy. In this work we characterize the role of *HB21* in the control of the proliferative arrest associated with the end of flowering. We observed that *HB21*, together with *HB40* and *HB53*, accumulate in the inflorescence apexes at the end of flowering promoting the cessation of inflorescence meristem activity. We also show that *HB21* induction of in young apexes is sufficient to induce flower and meristem arrest, likely mediated by an increase in ABA responses. Thus, our work confirms the parallelism proposed between dormant meristems and the arrested meristem at the end of flowering, which appear to be regulated by common pathways, and propose ABA as a new regulator in the control of inflorescence meristem arrest.

## Introduction

The final number of flowers that an inflorescence produces is influenced by the time that the inflorescence meristem is active. Thus, the flowering period, understood as the time that an inflorescence is able to generate new flowers, has a strong effect on the final yield of seeds and fruits. The flowering period initiates with the floral transition, a highly regulated developmental process that integrates multiple signals, both endogenous and exogenous (Andrés and Coupland, 2012; Blümel et al., 2015; Kinoshita and Richter, 2020; Freytes et al., 2021). In the other hand, the end of the flowering period corresponds with the moment when the apical meristem arrests and flower production ceases. Despite its ecological and agronomical interest, the end of flowering is still a largely uncharacterized process, even though our knowledge on this topic is increasing rapidly in the last years, and recent work suggests that the end of flowering, similarly to the floral transition, is a complex developmental process (González-Suárez et al., 2020; Wang et al., 2023).

At the genetic level, the end of flowering is controlled by the FRUITFULL-APETALA2 (FUL-AP2) pathway, that has been proposed to integrate age dependent and other endogenous signals. The AP2 transcription factor promotes meristem activity maintaining the expression of *WUSCHEL* (*WUS*), a stem cell identity gene (Laux et al., 1996; Mayer et al., 1998; Wurschum et al., 2006; Zhao et al., 2007). The MADS-box transcription factor FUL promotes the end of flowering, in part, by the direct repression of the *AP2* gene and others member of the *AP2* clade (Balanzà et al., 2018). FUL represses *AP2* together with the miR172. Thus, while *ap2* mutant combinations with other mutants of the *AP2* family show an early end of flowering, the *AP2 mir172* resistant alleles cause a delayed meristem arrest and *ful* mutants do not cease meristem activity and are able to produce flowers until the death of the plant (Balanzà et al., 2018; Merelo et al., 2022).

In addition to this genetic control, the end of flowering is also regulated by hormone signaling. It has been described that auxin export from developing fruits can trigger the meristem arrest by mechanisms that are still unclear (Ware et al., 2020; Goetz et al., 2021). Cytokinins (CK) have been also related to the control of the end of flowering. It has been shown that before meristem arrest at the end of flowering the CK response decreases in the shoot apical meristem, disappearing at the moment of meristem arrest (Merelo et al., 2022; Walker et al., 2023). The decrease in CK response is correlated with a decrease in the cell division rate and with the levels of the meristem marker WUS (Merelo et al., 2022).

In addition to the described factors, seed development is a strong meristem arrest promoter. In sterile mutants, or in plants where flowers are removed continuously, the inflorescence meristem remains active for longer, ending the flowering period with the differentiation of the shoot apical meristem in a terminal floral structure (Hensel et al., 1994; Balanzà et al., 2019).

In general, the meristem arrest of the inflorescence at the end of flowering can be considered a kind of dormancy. Transcriptomic analysis of arrested meristems at the end of flowering show a high degree of similarity with dormant meristems, presenting low mitotic activity and the activation of responses related to stress and growth inhibitory hormones as the abscisic acid (ABA) (Wuest et al., 2016). In agreement with this, meristem arrest at the end of flowering can be reverted by seed/fruit removal, *AP2* induction or CK treatments (Hensel et al., 1994; Balanzà et al., 2018; Merelo et al., 2022). Bud dormancy is controlled by the TCP transcription factor *BRANCHED1* (*BRC1*), directing the cell growth arrest responses that avoid the activation of the axillary meristems (Aguilar-Martínez et al., 2007). ABA signaling is one of the growth arrest responses controlled by BRC1 through the activation of three homeodomain leucine zipper (HD-ZIP) transcription factors: *HOMEOBOX PROTEIN 21* (*HB21*), *HB53* and *HB40*. These three genes enhance the expression of the *9-CIS-EPOXICAROTENOID DIOXIGENASE 3* (*NCED3*), a key gene in the ABA biosynthesis (Iuchi et al., 2001; Tan et al., 2003), triggering ABA accumulation (González-Grandío et al., 2017). Recently we have shown that downstream the FUL-AP2 pathway that control the end of flowering process there are some genes involved in the control of bud dormancy. AP2 is a direct repressor of the *HB21*, and when *AP2* is induced in active inflorescence meristems, the expression of *HB21, HB53* and in a lesser extent, *HB40*, are repressed, together with the ABA responses associated to the end of flowering and meristem arrest (Yant et al., 2010; Martínez-Fernández et al., 2020).

In this work we have characterized the expression of *HB21* during inflorescence development, as well as the contribution of *HB21, HB53* and *HB40* during the meristem arrest induction at the end of flowering. Transcriptomic analysis also indicate that these genes induce similar responses in the inflorescence apexes than the observed in dormant axillary buds, pointing out ABA as a new player in the control of meristem arrest at the end of flowering.

## Results

### HB21 accumulates in the inflorescence apex close to the end of flowering

The role of *HB21*, together with *HB40* and *HB53*, in the maintenance of bud dormancy in Arabidopsis has been previously described in detail (González-Grandío et al., 2017). Interestingly, it has also been suggested that HB21 could be involved in the control of flowering termination participating in inflorescence meristem arrest, which has been proposed to be a type of meristem dormancy (Martínez-Fernández et al., 2020). However, the relevance of HB21 during this developmental process is still unclear.

To understand the role of HB21 during inflorescence meristem arrest we decided to study its expression pattern in the inflorescence apex during the entire flowering period using a *proHB21:GUS* reporter line (González-Grandío et al., 2017). The GUS signal was absent in the inflorescence apex one week after bolting (wab) (Fig. 1A), as well as at two wab (Fig. 1B). At three wab, when the proliferative capacity of inflorescence meristem declines (Merelo et al., 2022), the GUS signal started to be detected in the base of the flower primordia at the inflorescence apex (Fig. 1C), being evident at 4wab and when the meristem is arrested (Fig. 1D-E). Interestingly the GUS signal never was detected in the SAM itself (Fig. 1A-E)

**Figure 1:**
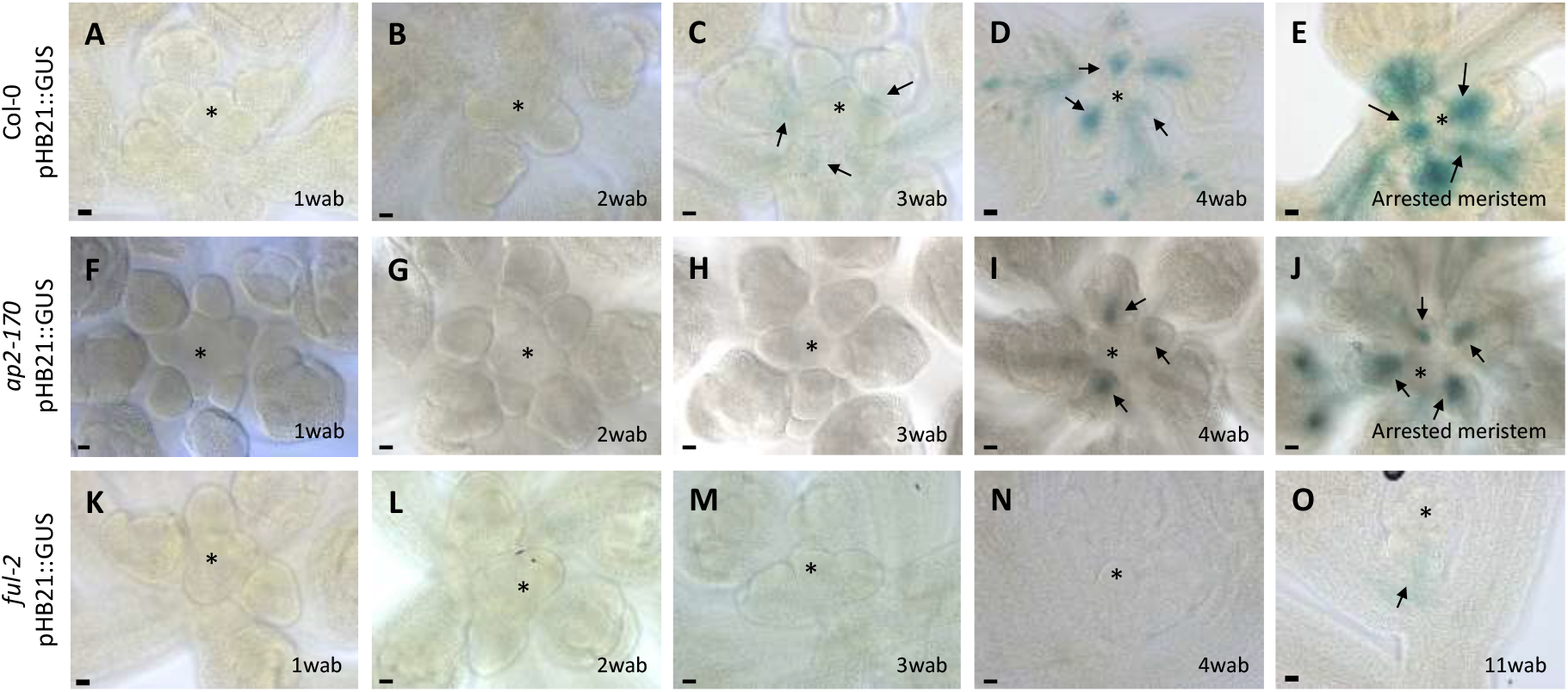
HB21 expression pattern accumulates close to the end of flowering. Histochemical detection of GUS activity driven by the HB21 promoter in Arabidopsis inflorescence apex. **(A-E)** Expression of pattern *proHB21:GUS* in WT background one wab (A), two wab (B), three wab at the onset of the low proliferative phase, when weak activity of the reporter is detected (C), four wab, when upregulation of the reporter is clear (D) and in arrested meristems, where the HB21 expression reaches a maximum (E). **(F-J)** Expression pattern of *proHB21:GUS* in the *ap2-170* background one wab (F), two wab (G), three wab (H), four wab, when the *HB21* expression begins (I) and in arrested meristems (J). **(K-O)** Expression pattern of *proHB21:GUS* in *ful-2* background one wab (K), two wab (L), three wab (M), four wab (N) and at eleven wab, where the plant is entering senescence, a slight HB21 expression can be detected (O). Arrowheads point to the floral primordia. Asterisk marks the SAM. Bar = 20μm.

To determine that the expression detected in the apex was related to the developmental arrest of the SAM and not a mere temporal correlation with inflorescence age, we decided to introduce the *proHB21:GUS* reporter in the *ap2-170* and *ful-2* mutant backgrounds, in which the activity of the inflorescence SAM is extended. *ap2-170* is a miR172 resistant allele that presents a delayed meristem arrest caused by the AP2 accumulation (Balanzà et al., 2018). It has been shown that AP2 is a direct negative regulator of *HB21* (Yant et al., 2010; Martínez-Fernández et al., 2020). The *proHB21:GUS* reporter in the *ap2-170* mutant was not detected during the first three wab (Fig1. F-H). The GUS signal started to be detected at 4wab (Fig.1 I), one week later than in the control line (Fig. 1C). 5wab, when meristem arrest is observed in the *ap2-170* mutant, the signal was evident in this line (Fig.1 J), in a similar pattern to the observed in the control line at 4 wab (Fig.1 D). This result suggest that the delayed meristem arrest observed in the *ap2-170* mutant could be associated with a delayed activation of *HB21* in the shoot apex.

In the *ful-2* mutant, the inflorescence meristem never experiences a full arrest (Merelo et al., 2022), partly due to the de-repression of several members of the *AP2* family, including *AP2* (Balanzà et al., 2018). In the *ful* mutant, the reporter signal was not detected during the entire flowering period (Fig. 1K-N), and only a weak GUS signal could be observed several weeks (11wab) after the onset of the arrest in wildtype plants, when the *ful* plants show conspicuous signs of overall senescence (Fig. 1O).

The inflorescence proliferative arrest is associated with a decline in WUS expression in the shoot apical meristem, where is no longer detected at the arrested stage (Balanzà et al., 2018; Merelo et al., 2022). To determine precisely when *HB21* accumulates in the shoot apex, we performed a simultaneous analysis of *WUS* and *HB21* expression patterns during inflorescence development by introducing a *pWUS::GFP:WUS* reporter into the *proHB21:GUS* line. Individual apices were sequentially processed for confocal detection of GFP:WUS and GUS analysis. As previously described, the *WUS* reporter was expressed at uniform levels until 3 wab, when it declined and eventually disappeared in arrested meristems at 4 wab (Fig. 2A). The decline in *WUS* expression at 3wab coincided with the initial expression of *HB21*, and the *WUS* switch-off a week later with the highest expression of *HB21* (Fig. 2B). The negative correlation between *WUS* and *HB21* supports again the idea that *HB21* initiates its expression when the shoot apical meristem enters the low proliferative phase that leads to flowering termination, reaching its maximum level when meristem arrests.

**Figure 2:**
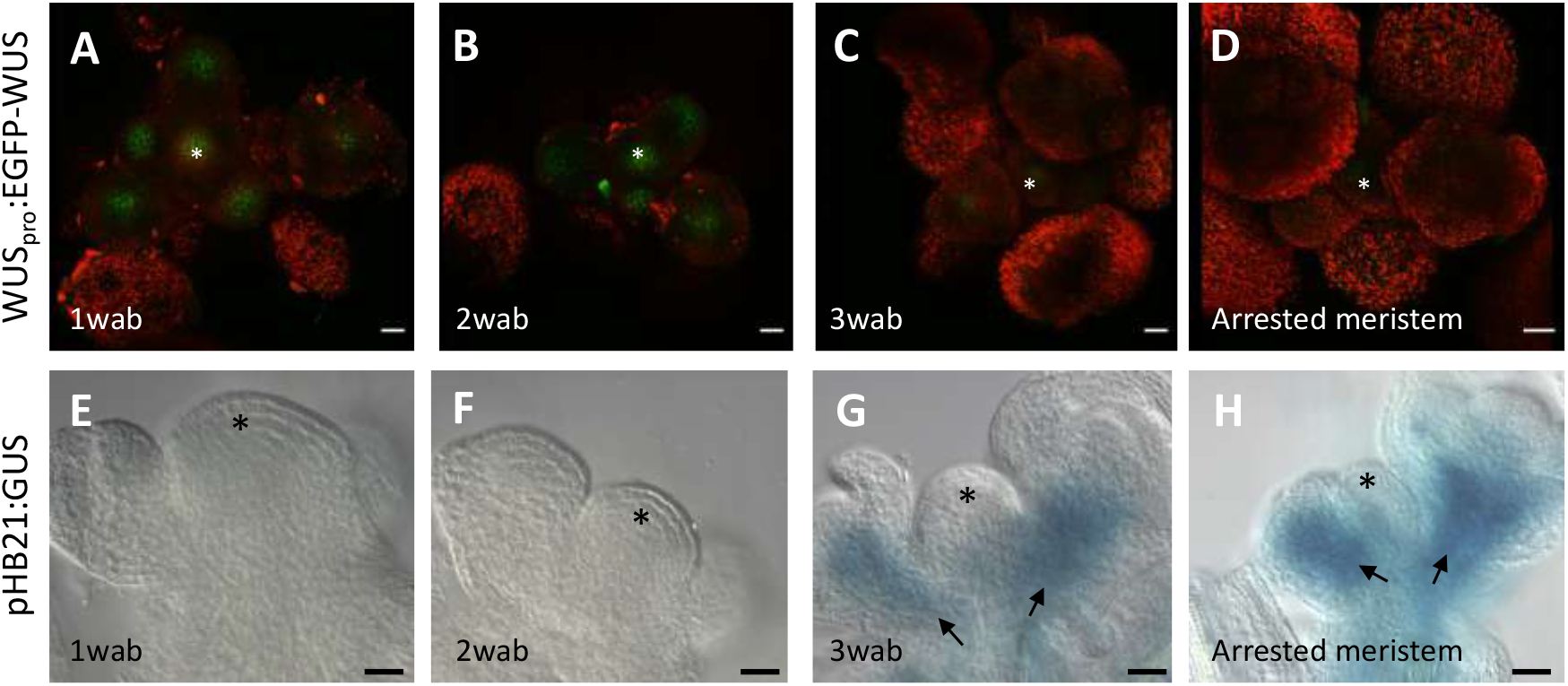
WUS accumulation negatively correlates with HB21 activation *pWUS::GFP:WUS* and *proHB21:GUS* lines were analyzed, the same confocal imaged apexes were sequentially processed for GUS analysis. **(A-D)** Expression of *pWUS::GFP:WUS* (green) one wab (A), two wab (B), three wab (C) and four wab (arrested) **(D-H)** Histochemical detection of GUS activity driven by the HB21 promoter at one wab (E), two wab (F), three wab (G) and arrested (H). At three wab WUS expression declines (C) coinciding with HB21 upregulation (G), and in arrested meristems WUS expression is no longer detected while HB21 promoter activity is at its highest (D, H). Arrowheads point to the floral primordia. Asterisk marks the SAM. Bar = 20 μm.

We also analyzed the signal of the *proHB21:GUS* reporter in the inflorescence apex after meristem reactivation by fruit removal (Hensel et al, 1994). Once the meristem starts to produce new flowers, no GUS signal was detected in the apex (Suppl. Fig. 1) reinforcing the idea that *HB21* expression associates with meristem arrest.

### *HB21* is able to induce meristem arrest when is expressed in the SAM

The negative correlation of inflorescence meristem activity and HB21 expression suggested that the HB21 protein could be triggering the dormant state associated to proliferative arrest at the end of the flowering period. To test this hypothesis, we generated an inducible line of *HB21* (p*35S:Lh:GR»HB21*) (Moore et al., 2006). For characterization, two independent lines were selected by their different response to Dexamethasone (Dex) treatment in leaves (line #7 (strong induction) and #21 (mild induction). We treated inflorescence main apexes 2 wab, a stage where meristem is fully active, with a drop of Dex or mock and checked the plants 5 days after treatment. The mock treated apexes continue with a normal development, opening new flowers associated to stem elongation, while the DEX treated apexes showed different phenotypes (Fig. 3A-C). In line #7 (mild) plants, Dex treatmemt induced flower primordia development arrest (Fig. 3B), one of the landmark events that occurs during the end of flowering process (Ref Benn?). However, induced line #7 plants did not show a visible effect in SAM activity, which stayed active producing new floral primordia. In the other hand, line #21 (strong) showed a clear response to treatment on the whole meristem apex, inducing flower primordia senescence and SAM arrest (Fig. 3C). This result indicates that HB21 is able to regulate the inflorescence development when expressed locally in the meristem apex, affecting flower development progression and meristem activity. Then, we combined the strong inducible *HB21* line #21 with the *ap2-170* and *ful-2* mutations, that cause a delay in meristem arrest and where expression of the endogenous *HB21* gene is repressed. In the *ap2-170* background, the induction of *HB21* at 2 wab generated two kinds of phenotype respect to the mock-treated plants (Fig. 3D-F). Some plants showed only the arrest of floral primordia development observed in the mild line #7 in the wild type background (Fig. 3B, 3E), while the rest were able to arrest both the growth of the SAM together with the senescence of the flowers (Fig. 3F), although the effect was less severe than the observed in line #21 induced plants in the wild type background (Fig. 3C). Interestingly, in the *ful-2* mutant, the induction of *HB21* did not affect inflorescence development, and five days after the DEX treatment the plants were indistinguishable from the mock-treated control plants (Fig. 3G-I. The results obtained with the *HB21* inducible line indicate that HB21 is sufficient to induce the arrest of the developing structures in the shoot apex, both flowers and meristem, and these phenotypes could be dependent on the HB21accumulation level, but also that FUL might be required for these functions.

**Figure 3:**
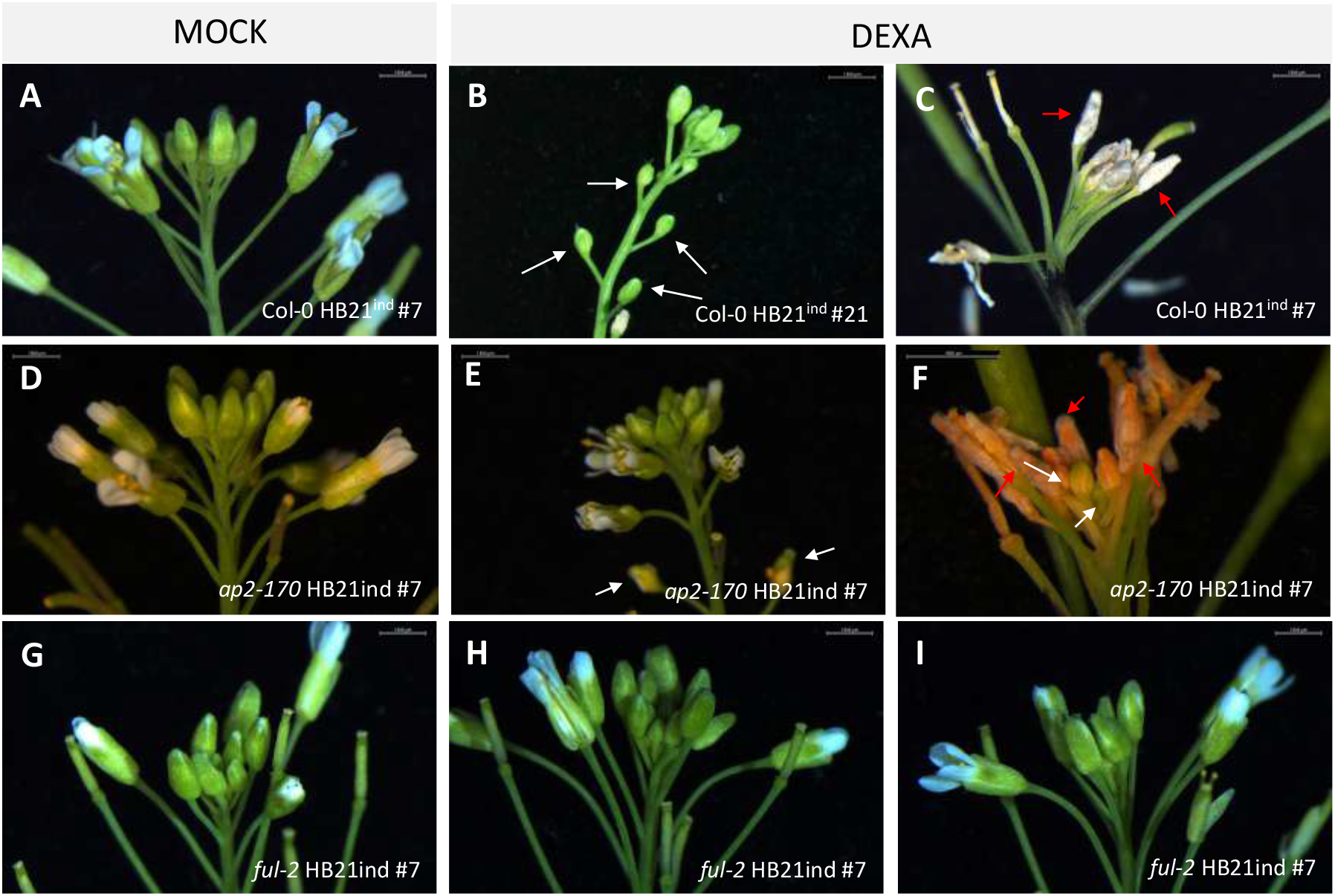
*HB21* induction induces flower and meristem arrest. Effect of the induction of *HB21* in proliferating inflorescences of WT, *ap2-170* and *ful-2* plants, 2wab. **(A)** WT *HB21*ind strong #7 line plant, mock-treated, show normal development **(B)** WT *HB21*ind mild line #21 plant after Dex treatment, where flower primordia are arrested. **(C)** WT *HB21*ind strong #7 line after Dex treatment, where arrested meristem and flower primordia senescence are observed. **(D)** *ap2-170 HB21*ind #7 mock-treated **(E)** *ap2-170 HB21*ind #7 show milder effects after Dex treatment, including flower primordia arrest. **(F)** *ap2-170 HB21*ind #7 where Dex treatments caused strong effects, including arrested inflorescence meristem and arrested senescing floral primordia. **(F-I)** *ful-2 HB21*ind #7 both mock-treated (F) and Dex-treated (H,I) showed no visible effects of the treatment. White arrows point to arrested flower primordia, red arrows point to senesced flower primordia, blue arrows point to reactivated flower primordia. Bar = 1,5mm.

### HB21 controls meristem arrest redundantly with HB53 and HB40

Transcripts of *HB21* accumulate at the end of flowering and its induction in the inflorescence apex is able to arrest the meristem activity. In agreement with this, it was previously proposed that *HB21* could modulate the end of flowering promoting meristem arrest based on the observation of *hb21-2* mutants (SAIL_790_D09.v1), which produced more flowers before inflorescence arrest than the control plants (Martínez-Fernández et al., 2020). To further characterize the role of *HB21* at the end of flowering we decided to check an additional allele, *hb21-1* (WiscDsLox468G4), previously described for its role in bud dormancy (González-Grandío et al., 2017). Both alleles are caused by the insertion of a T-DNA in the third exon of the gene (Suppl. Fig. 2A), in very close positions (*hb21-1* at position 8049808 and *hb21-2* at position 8049727). Surprisingly, both *hb21* alleles showed different phenotypes related with the end of flowering. Wild type plants produced 55,38 ± 3,73 fruits, while the *hb21-1* mutant produced 57,07 ± 7,15, in contrast with the 72,93 ± 6,10 fruits produced by the *hb21-2* mutant (Suppl. Fig. 2B). Because these alleles were likely not null, and to discard other putative second-site modifiers that could explain the disparity in phenotype of both mutant lines, we decided to generate additional mutant alleles by CRISPR/Cas9 genomic edition. We were able to generate plants with a deletion of 244 bp in the second exon of the gene, removing the HD domain. We named this new allele *hb21-3*,. These plants showed no apparent defects in plant architecture or organ development (Suppl. Fig. 3). When we compared the number of flowers produced before meristem arrest in the *hb21-3* mutant and the wild type plants a small, but significant difference was observed, producing 46.06 ± 2,95 and 41,83 ± 1,94 flowers respectively (Fig. 4A, 4B). This analysis indicates that the absence of *HB21* affects the number of flowers produced before arrest increasing the flower production around 10%.. This increase was milder in *hb21-3*, a null mutant allele, than in *hb21-2* (10 % increase vs 30% increase). This observation suggested that the allele *hb21-2* is not a real loss of function mutant, or that this mutant line could carry an additional insertion/mutation responsible of the observed phenotype.

**Figure 4:**
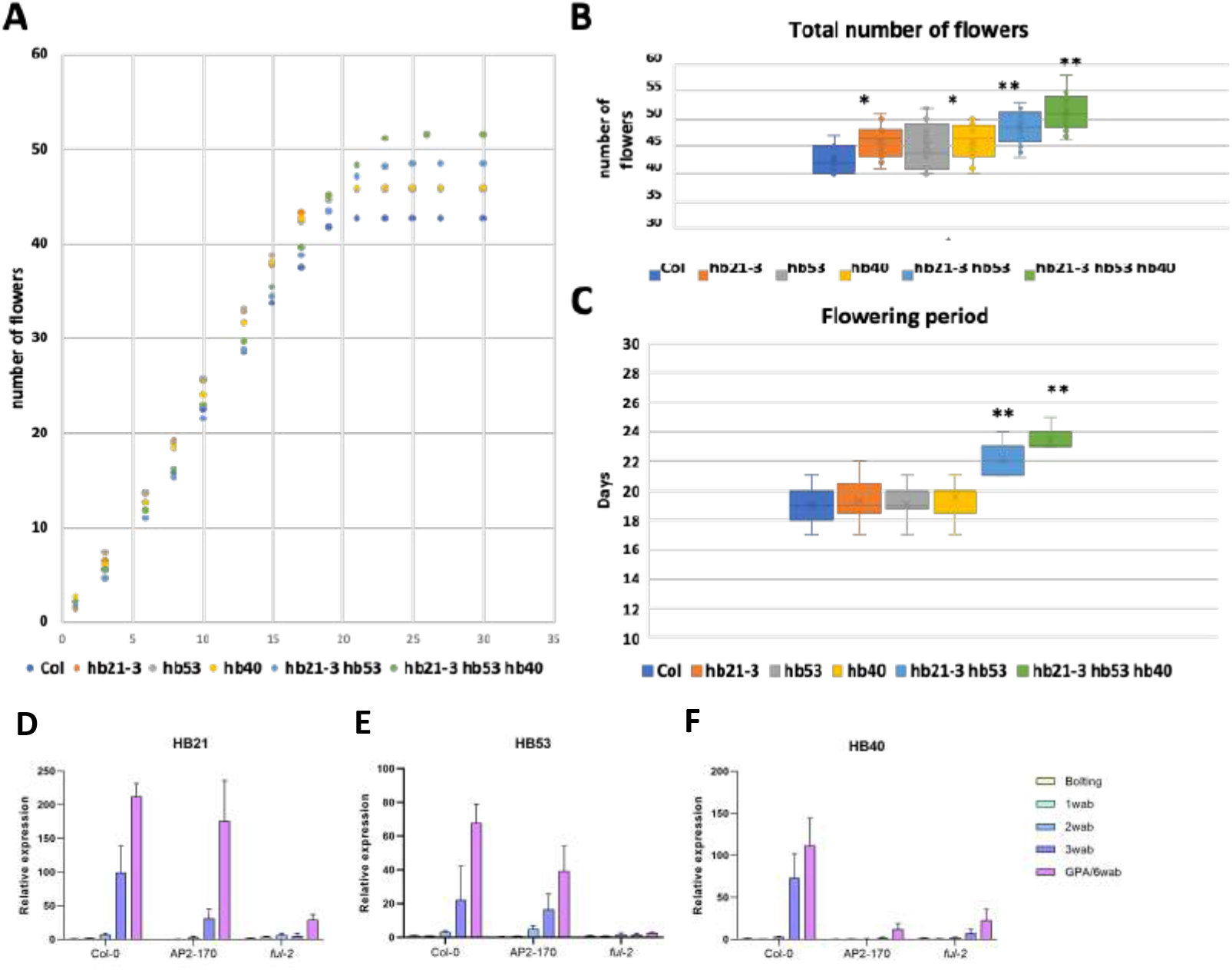
HB21, HB40 and HB53 act redundantly arresting meristem activity. **(A)** Number of flowers (accumulative) produced over time in WT, *hb21-3, hb53, hb40, hb21-3 hb50* and *hb21-3 hb53 hb40* backgrounds on the main inflorescence. **(B)** Total number of flowers produced in WT, *hb21-3, hb53, hb40, hb21-3 hb53* and *hb21-3 hb53 hb40* backgrounds on the main inflorescence, hb21-3 and hb40 produced more flowers than WT, and the combination *hb21-3 hb53* and *hb21-3 hb53 hb40* produced more flowers than the single mutants. **(C)** Duration of flowering in the main inflorescence in WT, *hb21-3, hb53, hb40, hb21-3 hb53* and *hb21-3 hb53 hb40* backgrounds, quantified as the interval between the first to the last flower in anthesis were observed **(D)** Transcript levels of *HB21* **(D)**, *HB53* **(E)** and *HB40* **(F)** at different time points of the flowering phase. All transcripts accumulate at the end of flowering in WT, in a reduced way in *ap2-170* mutants and are almost not detected in *ful-2* mutant.

Preliminary works with *HB21* indicate that this gene works redundantly with two additional genes, *HB40* and *HB53* in the control of bud dormancy (González-Grandío et al, 2017). *HB40* and *HB53* also accumulate at the end of flowering according to previously published transcriptomic data (Wuest et al., 2016), and *HB53* is repressed by AP2 induction in the inflorescence similarly to *HB21* (Martínez-Fernández et al., 2020). To assess whether HB21 could also work redundantly with HB40 and HB53 in the control of the end of flowering, we quantified the transcript accumulation of *HB21, HB53* and *HB40* during the reproductive phase in the inflorescence apex by qRT-PCR. For this purpose, we collected dissected inflorescence apexes from wild-type plants and *ap2-170* and *ful-2* mutants at the same time-points used in the *proHB21:GUS* analysis. The *HB21* transcript levels showed a clear increase at 4wab, when meristem arrest occurs, reaching its maximum level in apexes of 5wab (Fig. 4D). In the *ap2-170* mutant the expression pattern of *HB21* was similar to the observed in the wild type plants, but the expression levels at 4 and 5 wab were significantly lower than in the control (Fig. 4D). Finally, the *HB21* expression in the *ful-2* mutant was very low during all the inflorescence development, being only slightly up-regulated at the end of the inflorescence life, 11 wab (Fig. 4D). The changes in the expression levels detected by Q-PCR were in agreement with the GUS analysis reported for the *proHB21:GUS* line.

As observed for the *HB21* expression, *HB40* and *HB53* in the wild type plants started to accumulate in the inflorescence apex 3wab, reaching the highest expression level in arrested meristems (Fig. 4E, 5F). The expression of *HB40* in the *ap2-170* mutant, where meristem arrest is delayed, and the *ful-2* mutant, where meristem arrest never happens, was low in all time points assessed, accumulating slightly in the arrested *ap2-170* apexes (4wab) or in the apexes of *ful-2* mutants at 6wab (Fig. 4E). The expression of *HB53* in the *ap2-170* mutant was similar to the expression observed in the wild type plants, but was not upregulated at the same extent at the moment of meristem arrest (Fig. 4F). The levels of *HB53* in the *ful-2* mutant were always low (Fig. 4F). Our analysis confirms that *HB40* and *HB53* accumulate at high levels at the end of flowering as *HB21* does, and that their regulatory interaction with *AP2* and *FUL* are likely similar as well. Altogether, this supports the idea that the three HB genes might also act redundantly in the control of proliferative arrest at the end of the flowering phase.

To check this hypothesis, we characterized the *hb40-1* and *hb53-1* single mutants as well as the *hb21-3 hb53-1* and *hb21-3 hb53-1 hb40-1* double and triple mutants. The single mutants *hb53* and *hb40* produced in average a small increase in the final number of flowers (45,23 ± 4,32 and 45,83 ± 3,46) respect to the wild type control plants (41,83 ± 1,94) (Fig. 4A, 4B), but only the effect of *hb40* was statistically significant. The double mutant *hb21-3 hb53* produced a significant increase in the final number of flowers (47,69 ± 2,95) respect to the wild type plants, but not respect to the *hb21-3* (Fig. 4A, 4B). Finally, the triple *hb21-3 hb53 hb40* presented the stronger effect in the number of flowers produced by the inflorescence respect to the wild type plants as well as respect to the *hb21-3* single mutant, with a production of 49,25 ± 3,66 flowers.

The *HB* genes are expressed at the end of the flowering period, from the stage when the meristem declines in proliferative capacity until the arrest. Thus, *HB* genes should exert their function at this developmental stage. In agreement with this, the different mutants characterized did not affect meristem activity during the first weeks of flower production (Fig. 4A). We calculated then the flowering period of the inflorescence and no differences were observed between the wild type plants and all the single mutants (Fig. 4A, 4C). In contrast, both the double and triple mutant showed a clear extension in the flowering period respect to the wild type plants. Last opened flower in the wild type occurred at 19 ± 1,15 days after the opening of the first one (Fig. 4A, 4C). In the *hb21-3 hb53* and *hb21-3 hb53 hb40* mutants the last flower opened at 21,71 ± 0,99 and 21,81 ± 0,98 days respectively (Fig. 4A, 4C).

Our results indicated that the *HB21, HB53* and *HB40* genes regulate the final number of flowers produced in the inflorescence redundantly by extending the flowering period.

### HB21 controls meristem arrest controlling ABA response

Our results indicate that HB genes participate in the control of the end of flowering. To obtain more clues on the mechanism and the regulatory networks acting downstream these genes we decided to perform a whole transcriptomic analysis. Our inducible *HB21* line allow us to overcome the redundancy with *HB53* and *HB40*, so we decided to use this tool for this study. We treated inflorescence apexes of 2 wab *HB21* inducible lines (active meristems with low endogenous expression of *HB21*) with DEX or mock. Then, 6 hours post-treatment, we collected inflorescence apexes, removing all visible floral buds. 3 independent biological replicates for each treatment were used for RNA sequencing. Transcripts with a log2 fold change (FC) >1 and <-1, and a p-adjusted value < 0.05 were considered as differentially expressed genes (DEG) and selected for further analysis. We obtained 1143 DEG, 471 activated and 672 repressed by *HB21* induction (Suppl. Table 1)

With this list of DE genes we conducted a Gene Ontology (GO) analysis using the BiNGO tool (Maere et al., 2005) implemented for Cytoscape (Shannon et al., 2003), focusing in the enriched terms in the category Biological Process. For the downregulated DEGs we found 69 categories overrepresented, including the response to multiple stimulus and stress (Suppl. Table 2). Within the “response to stimulus” category, the response to hormones as jasmonic acid (12 genes), salicylic acid (11genes), abscisic acid (16 genes), auxins (15 genes) and ethylene (9 genes) stood out (Fig. 5A). The “response to stress” category included response to hypoxia (4 genes), to water deprivation (14 genes), to heat (14 genes), to cold (19 genes) and to oxidative stress (16 genes) (Fig. 5A). For the upregulated DEGs we found 53 categories overrepresented that included similar categories to the observed in the downregulated group (Suppl. Table 3), highlighting the response to abscisic acid (25 genes), water deprivation (31 genes) and cold (21 genes) (Fig. 5B). In addition, the response to light (17 genes) was also overrepresented together with the categories “leaf and organ senescence” (4 genes) (Fig. 5B). This analysis indicated that the induction of *HB21* is able to modulate the response to multiple stimulus, both endogenous and exogenous. It has been described that meristem arrest at the end of flowering is associated with an increased ABA response and resembles the state of bud dormancy. The transcription factor AP2 represses proliferative arrest, at least in part, by the repression of the ABA response. As AP2 is a direct negative regulator of *HB21* we decided to analyze which part of the role of AP2 in proliferative arrest is mediated by *HB21*. Thus, we compared the DEGs responding to the induction of *AP2* (Martínez-Fernández et al., 2020) with the DEGs responding to *HB21* induction. AP2 and HB21 exert opposite effects on meristem arrest: AP2 promotes meristem activity while HB21 promotes meristem arrest and senescence. Thus, we focused in genes that showed an opposite behavior in both experiments. We found that a total of 116 genes showed this pattern, 81 were up-regulated by HB21 and down-regulated by AP2 and 35 were down-regulated by HB21 and up-regulated by AP2 (Fig. 5C). We did a new GO analysis with these two subsets of genes (Suppl. Table 4) and we found that the group of genes up-regulated by HB21 and down-regulated by AP2 were also enriched the categories of response to stress, including the response to cold (8 genes) and the response to water deprivation (13 genes), and the response to endogenous stimulus, standing out the response to abscisic acid (8 genes) (Fig. 5D). In the complementary group, only the categories sulfur metabolic process (4 genes) and sulfate assimilation (3 genes) stood out. Our analysis suggested that *HB21* could mediate the ABA responses that appeared repressed by AP2. Once HB21 accumulates in the inflorescence apex, it could trigger the ABA response. In agreement with this hypothesis we identified in the list of up-regulated DEGs two key genes in the biosynthetic ABA pathway, *NCED3* and *NCED4* (Iuchi et al., 2001; Tan et al., 2003), with log_2_fold change values of 4,35 and 1,77 respectively. Thus, we hypothesized that up-regulation of *HB21* induces ABA biosynthesis and the subsequent ABA response, that could mediate flower and meristem arrest.

**Figure 5:**
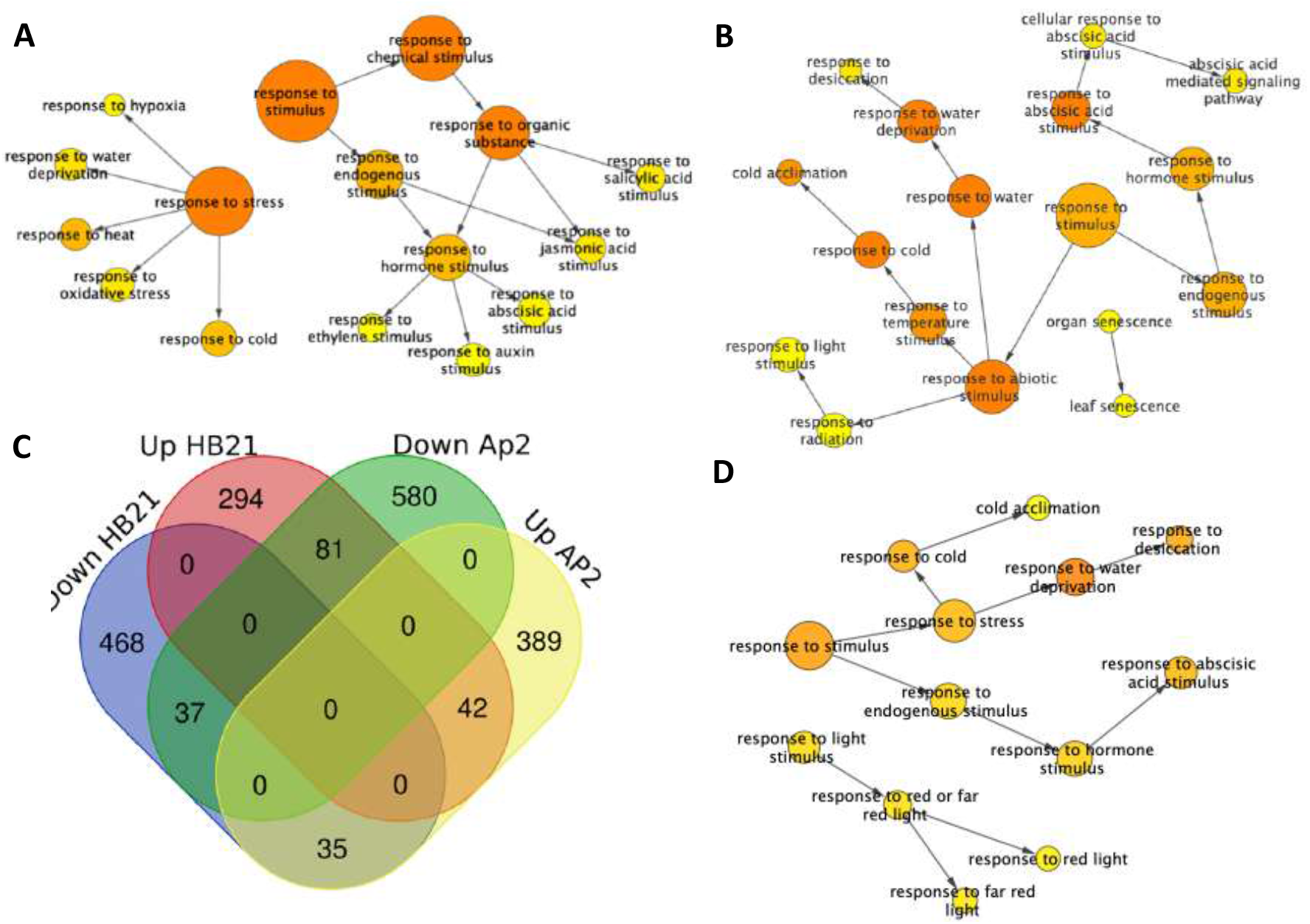
Functional enrichment analysis with overrepresented GO biological process categories. The analysis was performed for total DEG, upregulated **(A)** and downregulated **(B)** by *HB21* induction. The Venn diagram showed DEGs shared between *HB21* induction and *AP2* induction **(C)**. Go analysis of the 81 DEG up-regulated by *HB21* and down regulated by *AP2* **(D)**. All analysis share response to cold, water deprivation and abscisic acid stimulus (A,B,D). Circle size is proportional to gene numbers, and the color of each circle represents the enrichment Pvalue (hypergeometric test) for the GO term label on that circle, with orange representing the highest enrichment and yellow the lowest enrichment above the cutoff (Benjamin and Hochberg false discovery rate-corrected 0,05). Some categories were removed and the distance between noded was arranged manually to optimice readability. The figure and statistical analysis were generated using BiNGO software.

To test this hypothesis, we decided to apply local ABA treatments to inflorescence apexes of wild type plants and evaluate its impact on the meristem activity of wild type plants as well as of *hb21-3 hb53 hb40, ap2-170* and *ful-2* mutants. We applied a drop of a 70 μM ABA solution to inflorescence apexes during three consecutive days. The ABA treatment affected identically all the genotypes tested. While the control plants (mock treated) continued with a normal inflorescence growth (Fig. 6A, C, E, G), the ABA treated plants showed a clear reduction of growth. The ABA produced in the inflorescence apex an effect similar to the observed at the end of flowering, with a reduction of stem elongation, and the block of the development of the already formed flowers (Fig. 6B, D, F, H). Interestingly, all treated plants resume inflorescence growth after the cessation of the treatment, suggesting that continuous high levels of ABA are required to arrest the inflorescence progression.

**Figure 6:**
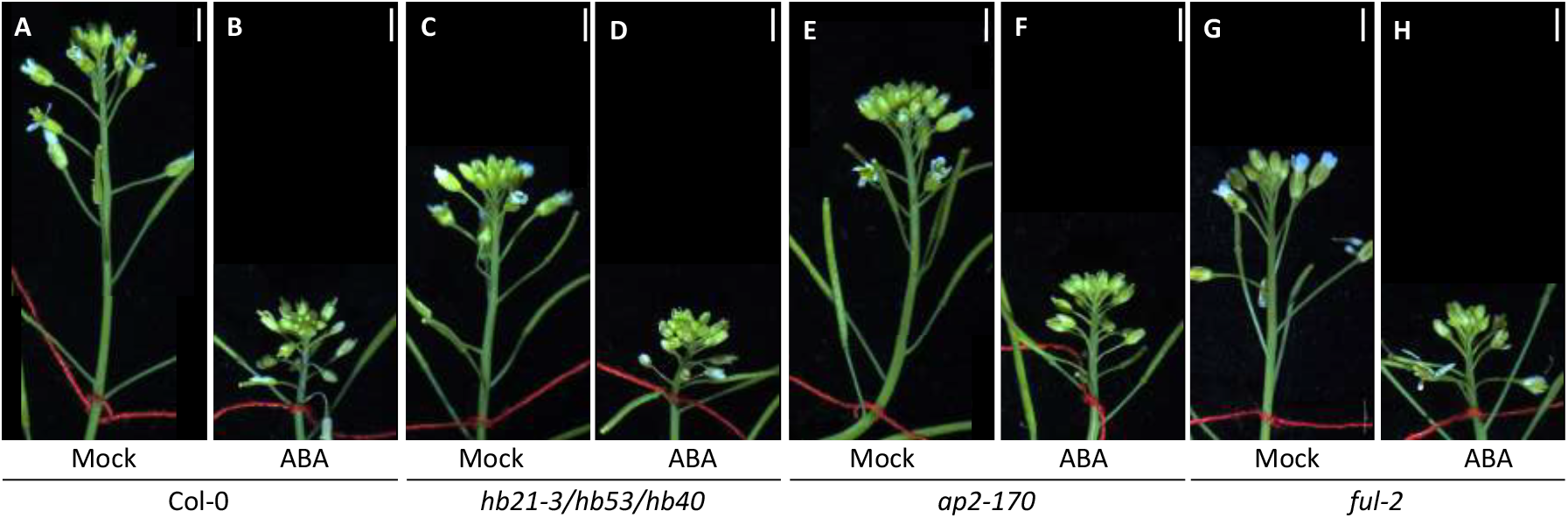
Abscisic acid arrests the development of the apical main inflorescence. ABA induction was performed on WT, *hb21-3/hb53/hb40, ap2-170* and *ful-2* backgrounds. WT’s apical inflorescence mock-treated control **(A)** and after treatment with ABA **(B)**, *hb21-3/hb53/hb40*’s apical inflorescence control **(C)** and after treatment with ABA **(D)**, *ap2-170*’s apical inflorescence control **(E)** and after treatment with ABA **(F)** and *ful-2*’s apical inflorescence control **(G)** and after treatment with ABA **(H)**. A decrease in growth is observed in all genotypes after the application of ABA. Bar=2.5mm.

## Discussion

During the last years, the mechanisms that control the developmental process of the end of flowering in monocarpic plants have started to be elucidated. It has been shown that the meristem arrest associated with the end of flowering is under genetic control, being the AP2 transcription factor a key negative regulator of the process (Balanzà et al., 2018; Martínez-Fernández et al., 2020). The meristem arrest could be affected by environmental factors such as light quality and photoperiod or temperature (Martínez-Fernández et al., 2020; González-Suárez et al., 2023) as well as by endogenous factors as auxins and CK hormones (Martínez-Fernández et al., 2020; Ware et al., 2020; Goetz et al., 2021; Merelo et al., 2022; Walker et al., 2023). In addition, transcriptomic analysis have also suggested that ABA signalling could mediate the meristem arrest (Wuest et al., 2016; Martínez-Fernández et al., 2020).

This work indicates that *HB21*, a direct target of AP2, together with *HB53* and *HB40*, could promote the ABA responses associated with the end of flowering through the activation of the ABA biosynthesis.

*HB21, HB53* and *HB40* has been described as part of the genetic regulatory network that control the dormancy of lateral buds in Arabidopsis. All three genes are activated by the TCP gene *BRANCHED1* in the axillary meristems. This activation occurs very early in development restricting the outgrowth of these meristems. We have shown that *HB21, HB53* and *HB40* are expressed in the inflorescence apex when the low proliferative phase of the meristem starts, preceding proliferative arrest, and at the same time that the levels of the meristem maintenance gene WUS begin to decline. Thus, as *HB21, HB53* and *HB40* do in axillary meristems, these genes could exert a similar function in the inflorescence apex, activating ABA biosynthesis and response. It has been described that at a similar time point for *HB* expression in the apex, the cytokinin response begins to decline (Merelo et al., 2022). CK are essential for meristem maintenance, promoting cell division and *WUS* expression (Bartrina et al., 2011; Schaller et al., 2014; Bartrina et al., 2017; Meng et al., 2017; Merelo et al., 2022). In the other hand, high ABA levels restrict growth and promote dormancy (Sreenivasulu et al., 2012; Reddy et al., 2013; Yao and Finlayson, 2015; González-Grandío et al., 2017).Thus, we could hypothesize that in the inflorescence apex, when *HB* genes start to be expressed, they cause *NCED* induction and subsequent ABA accumulation. It has been proposed that elevated ABA levels, as the induced by drought, could inhibit the biosynthesis of CK (Abe et al., 1997; Vaseva et al., 2008). Then the ABA response activated by *HB21* induction could participate in the progressive repression of the cytokinin response observed at the end of flowering. At the same time, the lower cytokinin levels could cause ABA hypersensitivity (Nishiyama et al., 2011), enhancing the ABA response. Then, ABA could be an important determinant in the control of meristem arrest. Supporting this hypothesis, in drought conditions, where ABA levels are elevated, an early and transitory inflorescence arrest has been described (Su et al., 2013). We have also confirmed that local ABA treatment in the inflorescence apex are able to arrest young inflorescences from different genetic backgrounds affected in the control of the end of flowering, suggesting that ABA is a final effector in the induction of inflorescence arrest.

This observation also explain the observed mild effects of our inducible HB21 line in the *ap2-170* background, where cytokinin response should be high due to the repression of the *KMD* genes (Kim et al., 2013; Martínez-Fernández et al., 2020). Interestingly, the induction of the same line in the *ful* mutant did not produce any effect in the inflorescence proliferative capacity, despite the ABA treatment was able to arrest the *ful* inflorescence. This suggests a clear dependence of the *HB21* effect on the presence of a functional FUL protein. Both genes, *HB21* and *FUL*, are meristem arrest promoters, but the expression of *HB21* depends on FUL. Once HB21 is present, it requires a functional FUL factor to affect meristem behavior. It could be interesting to asses if both proteins are able to interact, or if HB21 requires an additional factor that depends on FUL.

At the macroscopic level, the end of flowering is characterized by three different events: absence of stem elongation, developmental block of flower primordia and meristem arrest. After *HB21* induction in a young inflorescence apex, these three events are observed, suggesting that HB21 is sufficient to trigger the end of the flowering process. In the mild inducible line, the main phenotype observed was the developmental block of flower primordia, while in the strong line both stem elongation and meristem proliferation, in addition to the developmental block of flower primordia, were produced. These results suggest that HB21 accumulation could trigger the three events, but the requirement for each are different, being the flower developmental block the most sensitive. We have shown that in wild type plants, *HB21* is expressed mainly in the flower primordia. Then, we propose that HB21 should be acting initially on these tissues, inducing ABA synthesis and response, directing the flower primordia block. Once HB21 levels are high enough in these tissues, it could also contribute in the meristem arrest in a non-cell autonomous manner, well by the activation of a second factor, well by ABA transport to the adjacent tissues of the meristem (Kuromori et al., 2010). However, we can not discard that *HB21* could be acting directly on the meristem, as the transcript of this gene have been detected at high levels at the meristem arrest stage by RNAseq from micro-dissected samples of the shoot apical meristem (Wuest et al., 2016). These observations are also in agreement with the recently proposed progression of inflorescence arrest, in two independent and separate processes, the flowers developmental block and the shoot apical meristem arrest (Walker et al., 2023).

## Materials and methods

### Plant material and growth conditions

Arabidopsis thaliana seeds where stratified for 3d at 4ºC after sowing. Plants plants were grown in cabinets at 21 °C under LD (16 h light/8h dark) conditions, in a 2:1:1 (v/v/v) mixture of peat:perlite:vermiculite. All mutant plants and marker lines used in this study were in the Columbia background, except for *pWUS::GFP:WUS* which was in Landsberg. Mutant alleles and transgenic lines have been previously described: *ful-2, ap2-170, hb21-1, hb21-2, hb40-1, hb53-1*., *proHB21:GUS, pWUS::GFP:WUS*. The *35S::LhG4:GR»HB21* construct was generated by Gateway cloning of the *HB21* CDS from the REGIA collection (Paz-Ares and Consortium, 2002) into the pOpOn2.1 binary vector derived from the pOpOff2 vector (Moore et al., 2006). The mutant combinations were archived by parental crossing. The *hb21-3* allele was created by CRISPR method previously published (Wang et al., 2015) using the webtool (http://crispr.hzau.edu.cn/cgi-bin/CRISPR2/CRISPR) to design the two RNAg for generate a deletion. The deletion was checked by PCR and sequencing. Primer sequences used are detailed in Supplemental Table 5. In all cases, Arabidopsis was transformed with Agrobacterium tumefaciens strain C58 pM090 using the floral dip protocol (Clough and Bent, 1998), and both homocygous CRISPR lines and transgenic lines carrying a single transgene insertion were selected.

### β-Glucuronidase staining

For β-Glucuronidase (GUS) histochemical detection, samples were treated for 20 min in 90% ice-cold acetone and then washed for 5 min with washing buffer (50mM sodium phosphate pH7, 2mM ferrocyanide, 2mM ferricyanide, and 0,2% Triton X-100) and incubated O/N at 37 °C with staining buffer (washing buffer + 2mM X-Gluc). Following staining, plant material was fixed and cleared in chloral hydrate. Samples were mounted to be viewed under bright-field microscopy Leica DM5000.

### Confocal microscopy

Live imaging analyses were performed on a Stellaris 8 FALCON confocal microscope (Leica) using a water-dipping 40X objective. Reproductive shoot apices were imaged under water on MS medium plates, and with the stem embedded in the MS medium. To allow a proper exposition of the shoot apex during live imaging, all flower buds were carefully removed with clean tweezers and a fine needle. GFP was imaged using an argon laser emitting at the wavelength of 488 nm together with a 499-527 nm collection. Z stacks were acquired with a resolution of 8-bit depth, section spacing of 0.1 mm. More than five shoot apical meristems were observed.

### Dex treatment

Plants were grown in soil until 2 weeks after bolting. The induction of 35S::LhG4:GR»HB21 in the shoot apex of transgenic plants was carried out by putting one drop (3μL) of a Dex solution (10 μM Dex and 0.015% [v/v] Silwet L-77) or a control solution with an equivalent concentration of Silwet L-77 (mock) in the shoot apical meristem. Plants were observed 5 days later. For the RNAseq, inflorescence apices were harvested and dissected to eliminate older buds 6h after induction, three biological replicates were sampled, each containing about 16 inflorescence meristems.

### Quantitative RT-PCR

Inflorescence meristems were trimmed to remove old buds. Three biological replicates were sampled, each containing 16 inflorescence meristems. RNA was extracted using the E.Z.N.A. Plant RNA Kit (Omega Bio-tek) and DNase treated with EZNA RNase-Free DNase I (Omega Bio-tek). RNA concentration and purity were verified using a NanoDrop Spectrophotometer ND-1000 (Thermo Scientific). cDNAs were synthesized from 800 ng of total RNA using random hexamers and SuperScript IV (Invitrogen). The RT-qPCR was performed in the QuantStudio 3 Real-Time PCR (Thermo Fisher) and used SyberGreen to monitor double-stranded DNA synthesis. The Ct value was obtained from an automatic threshold. Results were normalized to the expression of the TIP41 reference gene. The 2^-ΔCt^ was shown as relative expression level. Three technical replicates were performed for each sample. Primer sequences used are detailed in Supplemental Table 5.

### Fruit/flower number quantification

For final fruit quantification elongated fruits were quantified in the main inflorescence for at least ten plants of each genotype after meristem arrest. For the accumulative number of flowers produced by the inflorescence, fruits and flowers in anthesis were counted each two/tthree days in the main shoot. Unhealthy or phenotypically altered plants were discarded. Experiments were replicated independently twice, obtaining comparable results, although only one experiment is represented in each figure.

### RNAseq

RNA for RNA-seq was obtained with the RNeasy Plant Mini Kit (QUIAGEN), DNase included in the Kit. RNA integrity was determined according to RNA Integrity Number values using a Bioanalyzer Chip RNA 7500 series II (Agilent). The RNA-seq was performed by Novogene Company United, with 20M reads. For the bioinformatic analysis, reads were aligned to the reference genome of Arabidopsis available at the TAIR database (Lamesch et al., 2012)) using TopHat (Trapnell et al., 2009) and Bowtie (Langmead et al., 2009) software. The abundance estimation of the transcripts was performed using the RSEM package (Li and Dewey, 2011) and the differentially expressed transcripts (fragments per kilobase million value) were estimated using Cufflinks (Trapnell et al., 2010). The sequences from DEG were annotated through BLAST search against the TAIR database. DEGs were analyzed using BiNGO tool (Maere et al., 2005) implemented for Cytoscape (Shannon et al., 2003), focusing in the enriched terms in the category Biological Process.

### Venn Diagrams

Venn diagrams were generated using the web tool (https://molbiotools.com/listcompare.php).

### ABA treatment

Plants were grown as described above. One week after bolting, 70 mM abscisic acid (ABA) or a control treatment (MOCK) was applied. For this, one drop (3μl) of ABA solution (70mM, 0.015% Silvet L77) or MOCK (0.015% Silvet L77) was added to the apical meristem. The treatment was repeated for three days and the plants were observed three days later.

### Statistic analysis

Two-tailed Student’s t-test was performed whenever two groups were compared. Statistical significance was determined at P < 0.05 unless otherwise indicated.

## Funding

This work was supported by the Spanish Ministerio de Ciencia e Innovación (grant no. RTI2018-099239-B-I00) and Generalitat Valenciana (grant no. PROMETEU/2019/004) to C.F.. and by the Spanish Ministerio de Economia, Industria y Competitividad (BES 2016078834) to V. S-G

## Author contributions

V.B., C.F., and V. S-G. conceived the original research plans; C.F. and V.B. supervised the experiments; V. S-G. performed most of the experiments; V. S-G. and V.B. analyzed the data; C.F. and V.B. wrote the article with contributions of all the authors.

## Acknowledgement

We thank Pilar Cubas for kindly provide the *hb21-1, hb40 and hb53* mutants and the reporter lines, and Javier Forment for help with the RNA-seq analysis.

## Notes

### Competing Interest Statement

The authors have declared no competing interest.

## References

Abe H, Yamaguchi-Shinozaki K, Urao T, Iwasaki T, Hosokawa D, Shinozaki K (1997) Role of arabidopsis MYC and MYB homologs in drought- and abscisic acid-regulated gene expression. The Plant Cell 9: 1859–1868

Aguilar-Martínez JA, Poza-Carrión C, Cubas P (2007) Arabidopsis BRANCHED1 Acts as an Integrator of Branching Signals within Axillary Buds. Plant Cell 19: 458–472

Andrés F, Coupland G (2012) The genetic basis of flowering responses to seasonal cues. Nat Rev Genet 13: 627–639

Balanzà V, Martinez-Fernandez I, Sato S, Yanofsky MF, Ferrandiz C (2019) Inflorescence Meristem Fate Is Dependent on Seed Development and FRUITFULL in Arabidopsis thaliana. Front Plant Sci 10: 1622

Balanzà V, Martínez-Fernández I, Sato S, Yanofsky MF, Kaufmann K, Angenent GC, Bemer M, Ferrándiz C (2018) Genetic control of meristem arrest and life span in Arabidopsis by a FRUITFULL-APETALA2 pathway. Nat Commun 9: 565

Bartrina I, Jensen H, Novak O, Strnad M, Werner T, Schmulling T (2017) Gain-of-Function Mutants of the Cytokinin Receptors AHK2 and AHK3 Regulate Plant Organ Size, Flowering Time and Plant Longevity. Plant Physiol 173: 1783–1797

Bartrina I, Otto E, Strnad M, Werner T, Schmulling T (2011) Cytokinin regulates the activity of reproductive meristems, flower organ size, ovule formation, and thus seed yield in Arabidopsis thaliana. Plant Cell 23: 69–80

Blümel M, Dally N, Jung C (2015) Flowering time regulation in crops—what did we learn from Arabidopsis? Current Opinion in Biotechnology 32: 121–129

Freytes SN, Canelo M, Cerdán PD (2021) Regulation of Flowering Time: When and Where? Current Opinion in Plant Biology 63: 102049

Goetz M, Rabinovich M, Smith HM (2021) The role of auxin and sugar signaling in dominance inhibition of inflorescence growth by fruit load. Plant Physiology 187: 1189–1201

González-Grandío E, Pajoro A, Franco-Zorrilla JM, Tarancón C, Immink RGH, Cubas P (2017) Abscisic acid signaling is controlled by a BRANCHED1/HD-ZIP I cascade in Arabidopsis axillary buds. Proceedings of the National Academy of Sciences 114: E245–E254

González-Suárez P, Walker CH, Bennett T (2020) Bloom and bust: understanding the nature and regulation of the end of flowering. Current Opinion in Plant Biology 57: 24–30

González-Suárez P, Walker CH, Bennett T (2023) FLOWERING LOCUS T mediates photo-thermal timing of inflorescence meristem arrest in Arabidopsis thaliana. Plant Physiology kiad163

Hensel LL, Nelson MA, Richmond TA, Bleecker AB (1994) The fate of inflorescence meristems is controlled by developing fruits in Arabidopsis. Plant Physiol 106: 863–76

Iuchi S, Kobayashi M, Taji T, Naramoto M, Seki M, Kato T, Tabata S, Kakubari Y, Yamaguchi-Shinozaki K, Shinozaki K (2001) Regulation of drought tolerance by gene manipulation of 9-cis-epoxycarotenoid dioxygenase, a key enzyme in abscisic acid biosynthesis in Arabidopsis. Plant J 27: 325–33

Kim HJ, Chiang YH, Kieber JJ, Schaller GE (2013) SCF(KMD) controls cytokinin signaling by regulating the degradation of type-B response regulators. Proc Natl Acad Sci U S A 110: 10028–33

Kinoshita A, Richter R (2020) Genetic and molecular basis of floral induction in Arabidopsis thaliana. Journal of Experimental Botany eraa057

Kuromori T, Miyaji T, Yabuuchi H, Shimizu H, Sugimoto E, Kamiya A, Moriyama Y, Shinozaki K (2010) ABC transporter AtABCG25 is involved in abscisic acid transport and responses. Proceedings of the National Academy of Sciences 107: 2361–2366

Lamesch P, Berardini TZ, Li D, Swarbreck D, Wilks C, Sasidharan R, Muller R, Dreher K, Alexander DL, Garcia-Hernandez M, et al (2012) The Arabidopsis Information Resource (TAIR): improved gene annotation and new tools. Nucleic Acids Res 40: D1202–10

Langmead B, Trapnell C, Pop M, Salzberg SL (2009) Ultrafast and memory-efficient alignment of short DNA sequences to the human genome. Genome Biol 10: R25

Laux T, Mayer KF, Berger J, Jurgens G (1996) The WUSCHEL gene is required for shoot and floral meristem integrity in Arabidopsis. Development 122: 87–96

Li B, Dewey CN (2011) RSEM: accurate transcript quantification from RNA-Seq data with or without a reference genome. BMC Bioinformatics 12: 323

Maere S, Heymans K, Kuiper M (2005) BiNGO: a Cytoscape plugin to assess overrepresentation of gene ontology categories in biological networks. Bioinformatics 21: 3448–9

Martínez-Fernández I, Menezes de Moura S, Alves-Ferreira M, Ferrándiz C, Balanzà V (2020) Identification of Players Controlling Meristem Arrest Downstream of the FRUITFULL-APETALA2 Pathway. Plant Physiology 184: 945–959

Mayer KF, Schoof H, Haecker A, Lenhard M, Jurgens G, Laux T (1998) Role of WUSCHEL in regulating stem cell fate in the Arabidopsis shoot meristem. Cell 95: 805–15

Meng WJ, Cheng ZJ, Sang YL, Zhang MM, Rong XF, Wang ZW, Tang YY, Zhang XS (2017) Type-B ARABIDOPSIS RESPONSE REGULATORs Specify the Shoot Stem Cell Niche by Dual Regulation of <em>WUSCHEL</em>. The Plant Cell 29: 1357–1372

Merelo P, González-Cuadra I, Ferrándiz C (2022) A cellular analysis of meristem activity at the end of flowering points to cytokinin as a major regulator of proliferative arrest in Arabidopsis. Current Biology 32: 749–762.e3

Moore I, Samalova M, Kurup S (2006) Transactivated and chemically inducible gene expression in plants. The Plant Journal 45: 651–683

Nishiyama R, Watanabe Y, Fujita Y, Le DT, Kojima M, Werner T, Vankova R, Yamaguchi-Shinozaki K, Shinozaki K, Kakimoto T, et al (2011) Analysis of Cytokinin Mutants and Regulation of Cytokinin Metabolic Genes Reveals Important Regulatory Roles of Cytokinins in Drought, Salt and Abscisic Acid Responses, and Abscisic Acid Biosynthesis. The Plant Cell 23: 2169–2183

Paz-Ares J, Consortium TR (2002) REGIA, an EU project on functional genomics of transcription factors from Arabidopsis thaliana. Comparative and Functional Genomics 3: 102–108

Reddy SK, Holalu SV, Casal JJ, Finlayson SA (2013) Abscisic Acid Regulates Axillary Bud Outgrowth Responses to the Ratio of Red to Far-Red Light. PLANT PHYSIOLOGY 163: 1047–1058

Schaller GE, Street IH, Kieber JJ (2014) Cytokinin and the cell cycle. Current Opinion in Plant Biology 21: 7–15

Shannon P, Markiel A, Ozier O, Baliga NS, Wang JT, Ramage D, Amin N, Schwikowski B, Ideker T (2003) Cytoscape: a software environment for integrated models of biomolecular interaction networks. Genome Res 13: 2498–504

Sreenivasulu N, Harshavardhan VT, Govind G, Seiler C, Kohli A (2012) Contrapuntal role of ABA: Does it mediate stress tolerance or plant growth retardation under long-term drought stress? Gene 506: 265–273

Su Z, Ma X, Guo H, Sukiran NL, Guo B, Assmann SM, Ma H (2013) Flower Development under Drought Stress: Morphological and Transcriptomic Analyses Reveal Acute Responses and Long-Term Acclimation in Arabidopsis. The Plant Cell 25: 3785–3807

Tan B-C, Joseph LM, Deng W-T, Liu L, Li Q-B, Cline K, McCarty DR (2003) Molecular characterization of the Arabidopsis 9-cis epoxycarotenoid dioxygenase gene family. The Plant Journal 35: 44–56

Trapnell C, Pachter L, Salzberg SL (2009) TopHat: discovering splice junctions with RNA-Seq. Bioinformatics 25: 1105–11

Trapnell C, Williams BA, Pertea G, Mortazavi A, Kwan G, van Baren MJ, Salzberg SL, Wold BJ, Pachter L (2010) Transcript assembly and quantification by RNA-Seq reveals unannotated transcripts and isoform switching during cell differentiation. Nat Biotechnol 28: 511–5

Vaseva I, Todorova D, Malbeck J, Trávníčková A, Macháčková I (2008) Response of cytokinin pool and cytokinin oxidase/dehydrogenase activity to abscisic acid exhibits organ specificity in peas. Acta Physiol Plant 30: 151–155

Walker CH, Ware A, Šimura J, Ljung K, Wilson Z, Bennett T (2023) Cytokinin signaling regulates two-stage inflorescence arrest in Arabidopsis. Plant Physiology 191: 479–495

Wang Y, Shirakawa M, Ito T (2023) Arrest, Senescence and Death of Shoot Apical Stem Cells in Arabidopsis thaliana. Plant and Cell Physiology 64: 284–290

Wang Z-P, Xing H-L, Dong L, Zhang H-Y, Han C-Y, Wang X-C, Chen Q-J (2015) Egg cell-specific promoter-controlled CRISPR/Cas9 efficiently generates homozygous mutants for multiple target genes in Arabidopsis in a single generation. Genome Biol 16: 144

Ware A, Walker CH, Šimura J, González-Suárez P, Ljung K, Bishopp A, Wilson ZA, Bennett T (2020) Auxin export from proximal fruits drives arrest in temporally competent inflorescences. Nat Plants. doi: 10.1038/s41477-020-0661-z

Wuest SE, Philipp MA, Guthörl D, Schmid B, Grossniklaus U (2016) Seed Production Affects Maternal Growth and Senescence in Arabidopsis. Plant Physiol 171: 392–404

Wurschum T, Gross-Hardt R, Laux T (2006) APETALA2 regulates the stem cell niche in the Arabidopsis shoot meristem. Plant Cell 18: 295–307

Yant L, Mathieu J, Dinh TT, Ott F, Lanz C, Wollmann H, Chen X, Schmid M (2010) Orchestration of the Floral Transition and Floral Development in Arabidopsis by the Bifunctional Transcription Factor APETALA2. Plant Cell 22: 2156–2170

Yao C, Finlayson SA (2015) Abscisic Acid Is a General Negative Regulator of Arabidopsis Axillary Bud Growth. Plant Physiol 169: 611–26

Zhao L, Kim Y, Dinh TT, Chen X (2007) miR172 regulates stem cell fate and defines the inner boundary of APETALA3 and PISTILLATA expression domain in Arabidopsis floral meristems. Plant J 51: 840–9

